# The challenge of RNA branching prediction: a parametric analysis of multiloop initiation under thermodynamic optimization

**DOI:** 10.1101/2020.01.03.893891

**Authors:** Svetlana Poznanović, Fidel Barrera-Cruz, Anna Kirkpatrick, Matthew Ielusic, Christine Heitsch

## Abstract

Prediction of RNA base pairings yields insight into molecular structure, and therefore function. The most common methods predict an optimal structure under the standard thermodynamic model. One component of this model is the equation which governs the cost of branching, where three or more helical “arms” radiate out from a multiloop (also known as a junction). The multiloop initiation equation has three parameters; changing those values can significantly alter the predicted structure. We give a complete analysis of the prediction accuracy, stability, and robustness for all possible parameter combinations for a diverse set of tRNA sequences, and also for 5S rRNA. We find that the accuracy can often be substantially improved on a per sequence basis. However, simultaneous improvement within families, and most especially between families, remains a challenge.

## 1 Introduction

Knowing the intra-sequence base pairings of an RNA molecule is typically a crucial step in understanding its function [1, 2]. Towards this end, thermodynamic optimization prediction methods remain essential tools for RNA structural biology [3], even as the ribonomics field moves forward [4, 5, 6, 7, 8, 9, 10, 11, 12].

A set of pseudoknot-free, canonical base pairs for a single-stranded RNA sequence is called a secondary structure. Each base pair defines a substructure, such as a hairpin loop or a base pair stack. Our interest here are the substructures known as multiloops (or junctions), which have three or more helical “arms” branching off. The canonical example for such a multiloop is the central single-stranded region in a 4-armed tRNA secondary structure. Multiloops determine the molecular shape [13] yet are some of the most difficult substructures to predict correctly [14].

The most common prediction methods use dynamic programing to efficiently generate a minimum free energy (MFE) structure as output [15, 16, 17, 18]. The free energy change from the unpaired RNA sequence is approximated under the nearest neighbor thermodynamic model (NNTM). The model, and associated parameters, are available online through the Nearest Neighbor Database (NNDB) [19]. The Δ*G* of a secondary structure is the sum of its substructure NNTM values. Here we analyze the initiation score, intended to approximate the entropic penalty, given to a multiloop.

Multiloop stability under the NNTM is the sum of two types of free energy changes. There is an initiation term (generally unfavorable) and then the various (favorable) values for the “stacking” of adjacent single-stranded nucleotides on base pairs in the loop. The stacking energies are based on experimental measurements [20, 21], but the initation is a linear function, originally chosen [20] for computational expediency, in three (learned) parameters;

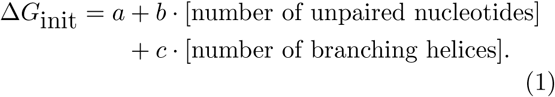

Previously, this simple entropy approximation was viewed with some concern [22, 23, 24], but recent results [25] demonstrate that it outperforms more complicated models in MFE prediction accuracy.

To achieve the full potential of this linear model for multiloop initiation, we should understand how MFE predictions depend on the (*a, b, c*) parameters. This is possible by applying mathematical theory to compute and analyze “RNA branching polytopes.” In this way, we can characterize the optimal branching of a given RNA sequence for every possible combination of (*a, b, c*). This approach, called a *parametric analysis*, permits us to quantify how much the accuracy can be improved, as well as other important characteristics like its stability and robustness.

We find that, on a per sequence basis, the accuracy can often be improved by a substantial amount, especially when it was originally low. However, the best predictions may require significantly different combinations of parameters. Hence, improving the average accuracy over a diverse set of sequences for a given RNA family, like tRNA or 5S rRNA, is much more challenging — but still possible.

However, our current approach cannot simultaneouly achieve this improvement for both the tRNA and 5S rRNA families tested. This result highlights that, while the linear model for multiloop initiation in Equation 1 can achieve very good accuracy, there may be a fundamental limit to possible improvements for MFE branching predictions.

## 2 Materials and Methods

We investigate how MFE prediction under the NNTM depends on multiloop initiation parameters. In our analysis, we vary the parameters (*a, b, c*) to characterize how the optimal branching changes, and its effect on important prediction characteristics.

As listed in Table 1, each major revision of the NNTM has changed the multiloop initiation parameters. The original “Turner89” parameters [20] are now no longer commonly used, but included here for completeness. The Turner99 ones [21] are still widely-used, as indicated by their listing in the NNDB [19]. The Turner04 multiloop model in the NNDB has a different form, but the recent study [25] showing the superior performance of Equation 1 reported using the values below.

**Table 1:** Multiloop initiation parameters over time.

| | $a$ | $b$ | $c$ |
| --- | --- | --- | --- |
| Turner89 | 4.6 | 0.4 | 0.1 |
| Turner99 | 3.4 | 0 | 0.4 |
| Turner04 | 9.3 | 0 | -0.6 |

Given an RNA sequence *R* as input, the MFE prediction algorithm has two parts. First, the minimum value, which is necessarily unique if (*a, b, c*) are fixed, is calculated. Next, at least one secondary structure *S* is computed whose free energy change Δ*G*_*S*_ is the MFE value. Often, only a single optimal structure is output, although it is possible to have two or more. The set of all MFE secondary structures can be computed by setting the free energy increment to 0 in the standard suboptimal structure algorithm [26].

### 2.1 Test sequences

Two families were tested: tRNA and 5S rRNA. Their native structures are well-characterized, and there are enough sequences available to generate a diverse test set. Computational limitations, discussed in Section 2.3.3, precluded a statistical analysis of longer sequences, like RNase P, at this point.

For each family, 50 sequences and their native base pairings were collected from the Comparative RNA Web (CRW) Site [27]. The pseudoknot-free secondary structures were used and only canonical base pairings are considered in accuracy calculations. The 50 sequences were arbitrary chosen so that their MFE prediction accuracies and GC content were distributed fairly evenly over the interval [0, 1]. Sequences, including accession numbers, length, MFE accuracy, and GC content, are listed in Supplementary Tables.

To assess the biological significance of their geometric characteristics, each set of biological branching polytopes was compared against two background distributions First, each test sequence was “shuffled” by the `ushuffle` program [28]. The new sequence has the same dinucleotide frequency [29, 30] as the original, but is otherwise randomized. Additionally, a set of uniformly random sequences, with the same length distribution as the original test set, was generated with a random number generator [31]. Each nucleotide has a 25% probability of being used in any given position.

### 2.2 Prediction characteristics

We evaluate the accuracy, stability, and robustness of the MFE predictions for different multiloop initiation parameter triples. We consider the three NNTM triples listed in Table 1 above, as well as three new triples, listed in Table 2 on page 6 which most improve predicction accuracy for the tRNA test sequences, for 5S, and for both families, respectively.

**Table 2:** Improved parameters from branching polytopes.

|  | Exact |  |  | Approximate |  |  |
| --- | --- | --- | --- | --- | --- | --- |
| | $a$ | $b$ | $c$ | $a$ | $b$ | $c$ |
| Best tRNA | 2729/250 | -53/375 | -261/100 | 10.9 | -0.1 | -2.6 |
| Best 5S | -52361/6160 | 789/3080 | 6873/1540 | -8.5 | 0.3 | 4.5 |
| Best both | 489/40 | 51/320 | -231/80 | 12.2 | 0.2 | -2.9 |

#### 2.2.1 Accuracy

Given the pseudoknot-free, canonical base pairings for a native secondary structure *S* and a corresponding MFE prediction *S*′ for that RNA sequence *R*, we score the accuracy as the *F*_1_-measure:

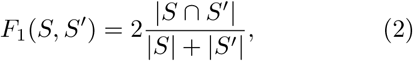

where |*S*| and |*S*′ | are the number of base pairs in *S* and *S*′, respectively, and |*S* ∩ *S*′ | is the number of true positive base pairs common to both structures. The minimum value 0 means no accurately predicted base pairs, while 1 means perfect prediction.

The accuracy of a multiloop initiation parameter triple for *R* is the average over all possible MFE secondary structures for that fixed (*a, b, c*). We report the average accuracy, with standard deviations, for each of the two RNA families tested.

### 2.2.2 Stability

The stability of a multiloop initiation triple is the amount those numbers can vary without changing the MFE prediction. Summary statistics are reported for each test family.

We first compute the amount each parameter can vary if the other two are fixed. This indicates the relative sensitivity of the MFE prediction to that parameter alone.

Next we consider how much the parameters can vary simultaneously. In particular, we investigate the rounding error effects, since the Turner parameters are calculated to 1 decimal precision. Hence, we consider a cube centered at a parameter triple, with edge lengths of .2, and compute the percentage of predictions which are stable within that cube. We repeat this for the (*a, c*) square with fixed *b* since we wish to understand the differences in sensitivity.

### 2.2.3 Robustness

Since we find that predictions typically have low stability, we also consider their robustness. The robustness of (*a, b, c*) will measure the similarity between predictions for other “nearby” parameter triples.

To compare two sets of MFE predictions, we calculate the worst best match between optimal structures for each parameter triple. More precisely, we compute the discrepancy as

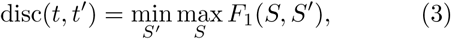

where *S* varies over all MFE secondary structures for *t* = (*a, b, c*), resp. *S*′ and *t*′ = (*a*′, *b*′, *c*′). Since the *F*_1_-measure is symmetric, it is used to score structural similarity here. Here 1 is two identical MFE secondary structures, and 0 is no common base pair.

The robustness of *t* over a distance *r* is then

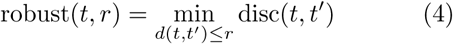

where the maximum metric

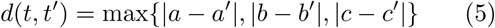

covers a cube with side lengths 2*r* centered at the parameter triple *t* = (*a, b, c*). We consider a progression of distances, starting with *r* = 0.1 and doubling every step until *r* = 3.2.

### 2.3 Parametric analysis

A description of the specific software used, as well as some background on the mathematical theory, have been published [32]. Here we focus only on the details relevant to this biological application.

#### 2.3.1 RNA branching signatures

The theory requires the thermodynamic optimization to be formulated as particular type of function, known as a *linear program*, in the parameters *a, b*, and *c*. Given the additive structure of the NNTM, this is easily done if we introduce a fourth “dummy” parameter *d*. Thus, the free energy change of a secondary structure *S* as a linear program in parameters (*a, b, c, d*) closely paralles Equation 1;

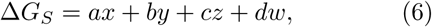

where

i. *x* is the number of multiloops in *S*,
ii. *y* is the total number of unpaired nucleotides in those multiloops,
iii. *z* is the total number of branching helices around those multiloops, and
iv. *w* is a remainder term which includes all other components of the Δ*G* calculation for *S* under the NNTM except those involving *a, b*, and *c*.

There is a crucial technicality, however. The set of base pairs *S* unambiguously determines Δ*G*_*S*_ — except for multiloop stabilities. Recall the second part of that calculation, often called the “dangling” energies, depends on the single-base stacking.

Since this information is essential to our parametric analysis, we work with *refined* secondary structures which, in addition to the usual base pairs, include the single-base stacking [32]. A refined secondary structure will be denoted 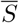.

For each 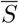, there is a single quadruple (*x, y, z, w*) which is its *branching signature*. However, there may be multiple refined secondary structures with the same signature. This is because, under the NNTM, it is possible to have different assignments of single-base stacking which leave *w* unchanged, or even different arrangments of base pairs which still yield the same (*x, y, z*) as well as *w*.

#### 2.3.2 RNA branching polytopes

Having formulated the thermodynamic optimization as a linear program, we can now — through the power of the mathematical theory — analyze the optimal branching signature for all of the (infinitely many) possible combinations of (*a, b, c*). To do this, we compute a geometric structure, know as the *branching polytope* for the given RNA sequence *R*.

For this *R*, there are only finitely many branching signatures possible, since there are only a finite number of possible refined secondary structures. The branching polytope is simply the smallest convex “envelope” enclosing these branching signatures.

A (filled) square is a 2-dimensional polytope, and a (solid) cube is one in 3d. In general, the “corners” of a polytope are called vertices and the flat sides are called faces; a cube has 8 vertices and 6 faces. Although an RNA branching polytope is significaly more complicated structurally than a cube, it is fundamentally the same type of mathematical object.

The mathematical theory says that if a linear program is optimized over a polytope, then the maximum and minimum are achieved on the boundary. (Intuitively, visualize sweeping a ruler across a square at a fixed angle to the horizontal.) Moreover, some combinations of parameters (that is, different angles of the ruler) give the same optimum while others give different ones.

The theory tells us that two different combinations of multiloop initiation parameters yield the same optimum branching signature if and only if they both lie in the same connected, convex region of the (*a, b, c, d*) parameter space. Hence, to understand the infinite parameter space, we “only” need to compute the finite number of optimal branching signatures on the boundary of the RNA branching polytope.

#### 2.3.3 Computional challenges

Although the number of branching signatures is finite for a given RNA sequence, it is far too large to compute the branching polytope directly from this set. Instead, we use the `pmfe` software [32] developed for this specialized purpose.

Critically, the software running time depends on the number of vertices plus the number of faces of the polytope being computed. RNA branching polytopes are structurally rather complex, and that complexity increases with sequence length. Hence, a tRNA computation takes about 2 hours, while 5S rRNA takes about a day; see Table 8 on page 9 for details.

**Table 3:** MFE prediction accuracy comparison.

| Parameters | tRNA |  | 5S rRNA |  |
| --- | --- | --- | --- | --- |
|  | Avg | Std | Avg | Std |
| Turner89 | 0.41 | 0.25 | 0.69 | 0.24 |
| Turner99 | 0.52 | 0.30 | 0.63 | 0.24 |
| Turner04 | 0.45 | 0.28 | 0.64 | 0.24 |
| Best tRNA | 0.75 | 0.24 | 0.50 | 0.21 |
| Best 5S | 0.36 | 0.19 | 0.74 | 0.19 |
| Best both | 0.73 | 0.26 | 0.61 | 0.22 |
| Best tRNA (approx) | 0.74 | 0.24 | 0.52 | 0.21 |
| Best 5S (approx) | 0.36 | 0.18 | 0.71 | 0.22 |
| Best both (approx) | 0.71 | 0.27 | 0.62 | 0.21 |

**Table 4:**
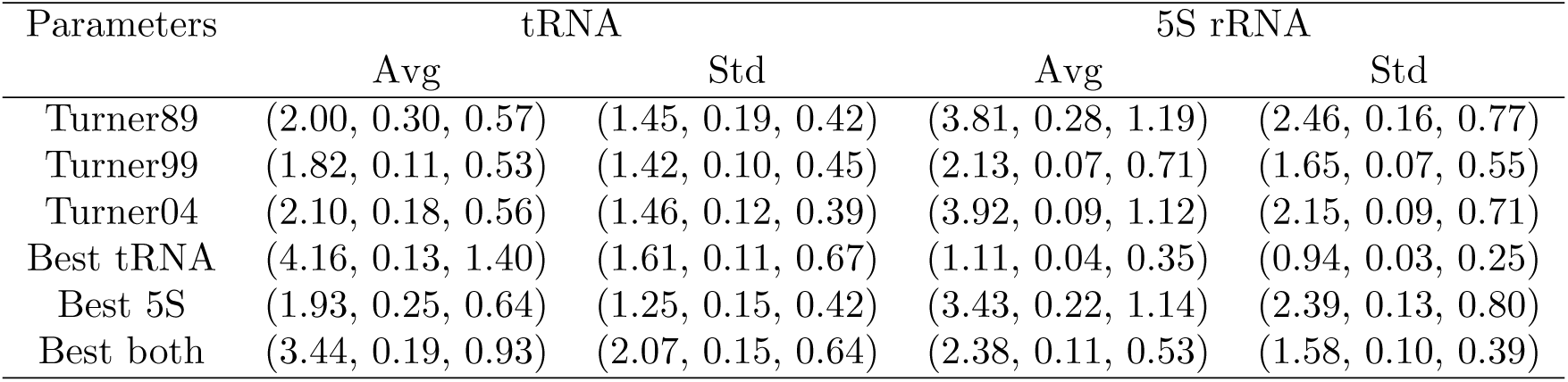
(*a, b, c*) stability for tRNA and 5S rRNA.

**Table 5:** Stability percentage for (*a, c*) under .1 rounding error.

|  | tRNA |  | 5S rRNA |  |
| --- | --- | --- | --- | --- |
| | fixed $b$ | $\Delta b \leq 0.1$ | fixed $b$ | $\Delta b \leq 0.1$ |
| Turner89 | 84 | 68 | 88 | 8 |
| Turner99 | 76 | 32 | 82 | 16 |
| Turner04 | 86 | 62 | 88 | 28 |
| Best tRNA | 98 | 58 | 78 | 6 |
| Best 5S | 86 | 70 | 88 | 76 |
| Best both | 88 | 62 | 82 | 40 |

**Table 6:** Robustness of tRNA prediction within error *r*.

| Parameters | $r = 0.1$ | | $r = 0.2$ | | $r = 0.4$ | | $r = 0.8$ | | $r = 1.6$ | | $r = 3.2$ | |
| --- | --- | --- | --- | --- | --- | --- | --- | --- | --- | --- | --- | --- |
|  | Avg | Std | Avg | Std | Avg | Std | Avg | Std | Avg | Std | Avg | Std |
| Turner89 | 0.84 | 0.30 | 0.71 | 0.34 | 0.47 | 0.27 | 0.33 | 0.19 | 0.11 | 0.13 | 0.03 | 0.04 |
| Turner99 | 0.71 | 0.31 | 0.55 | 0.29 | 0.41 | 0.21 | 0.26 | 0.18 | 0.07 | 0.09 | 0.03 | 0.04 |
| Turner04 | 0.77 | 0.31 | 0.58 | 0.31 | 0.38 | 0.20 | 0.24 | 0.19 | 0.06 | 0.08 | 0.03 | 0.04 |
| Best tRNA | 0.91 | 0.16 | 0.79 | 0.23 | 0.61 | 0.24 | 0.25 | 0.20 | 0.06 | 0.07 | 0.03 | 0.04 |
| Best 5S | 0.85 | 0.27 | 0.67 | 0.33 | 0.39 | 0.26 | 0.30 | 0.19 | 0.08 | 0.11 | 0.02 | 0.04 |
| Best both | 0.87 | 0.23 | 0.75 | 0.28 | 0.52 | 0.28 | 0.40 | 0.18 | 0.09 | 0.10 | 0.03 | 0.04 |

**Table 7:** Robustness of 5S rRNA prediction within error *r*.

| Parameters | $r = 0.1$ | | $r = 0.2$ | | $r = 0.4$ | | $r = 0.8$ | | $r = 1.6$ | | $r = 3.2$ | |
| --- | --- | --- | --- | --- | --- | --- | --- | --- | --- | --- | --- | --- |
|  | Avg | Std | Avg | Std | Avg | Std | Avg | Std | Avg | Std | Avg | Std |
| Turner89 | 0.91 | 0.24 | 0.86 | 0.28 | 0.65 | 0.34 | 0.36 | 0.22 | 0.11 | 0.08 | 0.03 | 0.03 |
| Turner99 | 0.78 | 0.27 | 0.61 | 0.28 | 0.45 | 0.22 | 0.29 | 0.17 | 0.08 | 0.06 | 0.04 | 0.03 |
| Turner04 | 0.82 | 0.29 | 0.74 | 0.30 | 0.44 | 0.24 | 0.30 | 0.17 | 0.08 | 0.06 | 0.04 | 0.03 |
| Best tRNA | 0.76 | 0.21 | 0.60 | 0.20 | 0.49 | 0.19 | 0.27 | 0.13 | 0.08 | 0.06 | 0.05 | 0.03 |
| Best 5S | 0.87 | 0.30 | 0.77 | 0.34 | 0.57 | 0.33 | 0.33 | 0.20 | 0.10 | 0.08 | 0.03 | 0.02 |
| Best both | 0.82 | 0.24 | 0.69 | 0.24 | 0.54 | 0.23 | 0.37 | 0.17 | 0.11 | 0.07 | 0.04 | 0.02 |

**Table 8:** Polytope computation time and structural complexity for tRNA and 5S rRNA.

| Family | # Seq | Length (nt) |  | Time (hr) |  | # Vertices |  | # Faces |  |
| --- | --- | --- | --- | --- | --- | --- | --- | --- | --- |
|  |  | Avg | Std | Avg | Std | Avg | Std | Avg | Std |
| tRNA | 50 | 74.38 | 1.89 | 1.86 | 0.26 | 703 | 56 | 2075 | 196 |
| shuffled | 50 | 74.38 | 1.89 | 2.05 | 0.35 | 718 | 73 | 2093 | 220 |
| uniform | 50 | 74.38 | 1.89 | 1.96 | 0.32 | 708 | 63 | 2072 | 213 |
| 5S | 50 | 121.38 | 3.62 | 23.15 | 3.33 | 2639 | 183 | 7649 | 524 |
| shuffled | 50 | 121.38 | 3.62 | 23.62 | 4.74 | 2606 | 251 | 7436 | 745 |
| uniform | 50 | 121.38 | 3.62 | 22.84 | 4.03 | 2523 | 228 | 7262 | 681 |

Increasing the sequence length by another 50 nucleotides (nt) to 175 increases the time to a week. The longest computation attempted thus far, for an RNase P sequence of length 354 nt, took more than 2 months. Hence, extending this analysis to more, longer sequences will require new algorithmic approaches to computing RNA branching polytopes.

#### 2.3.4 Data analysis

To generate the data reported, the RNA branching polytopes produced by `pmfe` were analyzed using the mathematical software `sage` [33]. Since multiple comparisons among summary statistics (averages and standard deviations) were often made, unless otherwise indicated, statistical significance of differences was assessed using a standard one-way analysis of variation (ANOVA) followed by Tukey Honestly Significant Difference (HSD) post-hoc tests at the 95% confidence level.

#### 2.3.5 Computing best accuracies

For each vertex of a branching polytope/region of the parameter space, we computed the prediction accuracy as the average over all refined secondary structures which attain the common MFE value. This identifies the most accurate parameters for that sequence.

However, to find the best prediction for a set of sequences involves considering the intersection of parameter regions for difference polytopes. It is computationally infeasible to consider all possibilities for our test families, so we restricted to searching for large subsets which achieve their best accuracy simultaneously. This is possible using an graph algorithm, implemented in `sage`, that searches for large “cliques.”

For this purpose, we built a graph for tRNA and one for 5S rRNA. Each graph vertex represents one of the 50 test sequences, and two vertices are conntected by an edge if their maximum accuracy can be achieved simultaneously. A clique is a subset of vertices where all possible edges occur in the graph. In this case, a large clique is a useful set of test sequences whose best parameters have nonempty intersection.

## 3 Results and Discussion

We address first the biological implications of our analysis, and defer the geometric details until later.

### 3.1 Biological implications

The explicit construction of 3d parameter decomposition for each RNA sequence enables us to determine how well the NNTM approximates the native secondary structure.

#### 3.1.1 Improving accuracy

We found that 89% of tRNA and 90% of 5S rRNA predictions can be improved. Figure 1 shows maximum accuracy per sequence over the Turner99 baseline, as well as average maximum over average baseline. However, it is not possible to achieve this maximum average for either test set because the common intersection for the best regions is empty.

**Figure 1:**
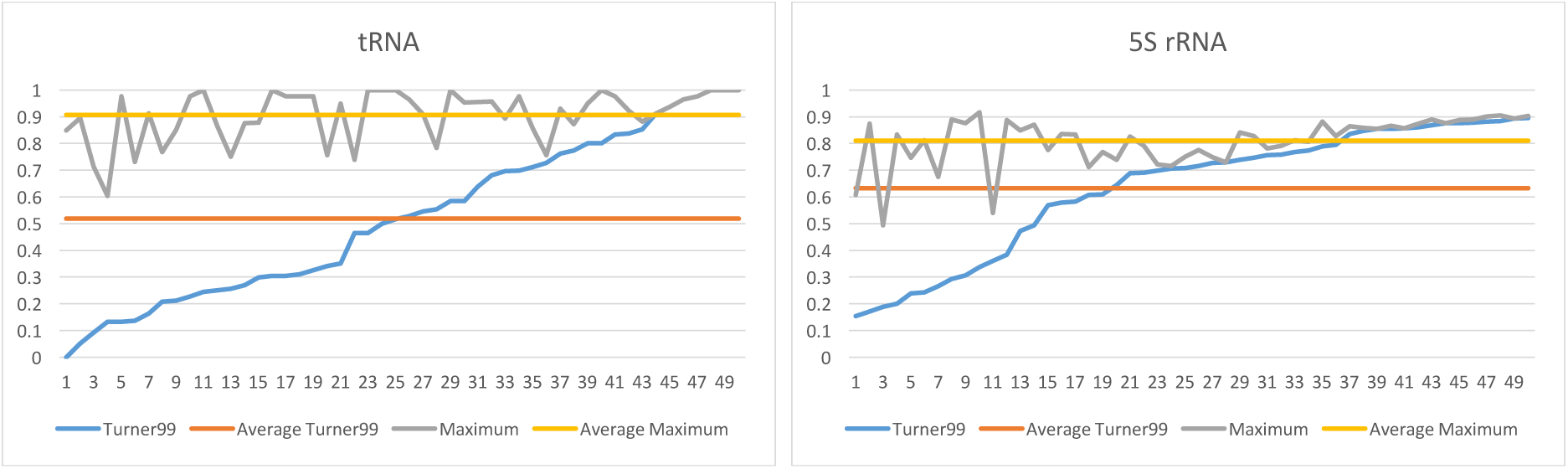
Per sequence improvements for tRNA and 5S rRNA test sets. Initial (Turner99) average accuracies are 0.52 (0.30) and 0.63 (0.24), resp. Most can be improved, by 0.39 (0.27) and 0.18 (0.21) on average, yielding maximum possible averages of 0.91 (0.10) for tRNA and 0.81 (0.09) for 5S rRNA. Differences within families have high statistical significance (*p* < 0.0002). Between family differences in initial accuracies are weakly significant (*p* = 0.0372) but maximum ones are not (*p* = 0.0701).

To find a better maximum average, we used the method in Section 2.3.5 which finds large subsets of sequences whose best accuracy is simultaneously achievable. We found 7 such subsets for tRNA and 4 for 5S rRNA. For each subset, we computed the common intersection of their best possible regions, and found its center. The prediction accuracies for the test sequences were then computed using this center as the multiloop initiation parameters.

The best (*a, b, c*) triples found for tRNA, for 5S rRNA, and for both are reported in Table 2. The first values listed are fractions because the software used computes over the rationals. To facilitate comparison with the current NNTM values, listed in Table 1 on page 2, the exact fractions were rounded to 1 decimal place, and are also reported in Table 2.

Table 3 lists average MFE prediction accuracy under these different parameter triples. Using the approximate values, rather than the exact fractions, can change the prediction; see Section 3.1.2 for further details. However, no significant differences between the exact and approximate prediction accuracies were identified.

Interestingly, the differences among the prediction accuracies within each family for the three Turner triples are not statistically significant either.

For tRNA, the differences between “best tRNA” and “best both” parameters were not statistically significant. However, the differences between these and the Turner parameters were, as well as the differences with the “best 5S.”

For 5S rRNA, only two types of significant differences were found: between “best tRNA” and the three Turner parameters as well as the “best 5S” ones.

We conclude that the “best” parameters for one family substantially lower the prediction accuracy for the other. However, the “best both” parameters raise the tRNA prediction accuracy considerably without negatively affecting the 5S rRNA predictions relative to the Turner parameters.

Our analysis here supports recent NNTM developments; the Turner04 parameters have a much larger *a* penalty than previously, but the *c* value is actually negative (so weakly favorable). The “best both” parameters have an even larger *a* penalty (12.2 versus 9.3) and a much more strongly favorable *c* value (−2.9 versus −0.6). Additionally, there is a small loop size penalty of *b* = 0.2.

#### 3.1.2 Stability

Table 4 reports distance to the closet region boundary in each dimension if the other two are fixed. Every (*a, b, c*) triple tested is most sensitive to changes in *b*, which weights the number of single-stranded nucleotides in a multiloop. This is because the regions overall are much thinner in this direction; see Table 10 and discussion of the general geometry of the parameter space decomposition in Section 3.2.2.

**Table 9:** Number of regions in (*a, b, c, d*) parameter space under constraints.

| Family | $d = 1$ | | bnd $d = 1$ | | $(b, d) = (0, 1)$ | | bnd $(b, d) = (0, 1)$ | |
| --- | --- | --- | --- | --- | --- | --- | --- | --- |
|  | Avg | Std | Avg | Std | Avg | Std | Avg | Std |
| tRNA | 517 | 42 | 320 | 36 | 46 | 6 | 29 | 5 |
| shuffled | 536 | 60 | 335 | 46 | 48 | 6 | 31 | 5 |
| uniform | 529 | 53 | 329 | 46 | 50 | 8 | 33 | 7 |
| 5S | 2109 | 164 | 1607 | 138 | 125 | 16 | 97 | 14 |
| shuffled | 2072 | 235 | 1564 | 201 | 127 | 21 | 98 | 19 |
| uniform | 2007 | 191 | 1515 | 165 | 128 | 20 | 100 | 18 |

**Table 10:** Average *d* = 1 bounded region dimensions in (*a, b, c*)

| Parameters | All | | $x > 1$ | |
| --- | --- | --- | --- | --- |
|  | Avg | Std | Avg | Std |
| tRNA | (26.80, 0.46, 2.55) | (3.15, 0.06, 0.49) | (5.07, 0.48, 2.63) | (1.04, 0.07, 0.54) |
| shuffled | (26.92, 0.49, 2.71) | (3.94, 0.09, 0.56) | (5.47, 0.51, 2.86) | (1.19, 0.09, 0.67) |
| uniform | (26.01, 0.48, 2.60) | (4.39, 0.08, 0.45) | (5.24, 0.50, 2.73) | (0.87, 0.09, 0.50) |
| 5S | (32.10, 0.41, 2.24) | (3.12, 0.04, 0.27) | (4.23, 0.42, 2.30) | (0.51, 0.05, 0.29) |
| shuffled | (27.26, 0.36, 1.92) | (3.76, 0.07, 0.37) | (3.75, 0.37, 1.97) | (0.65, 0.67, 0.38) |
| uniform | (27.06, 0.37, 2.00) | (3.62, 0.07, 0.37) | (3.92, 0.38, 2.07) | (0.65, 0.07, 0.39) |

Furthermore, each triple is least sensitive to *a*, whose stability is always at least 3 times *c*. Beyond this, no clear correlations were found.

We also consider perturbing the parameters simultaneously, within the 1 decimal rounding error. Table 5 gives the percentage of predictions which are stable when Δ*a, b, c* ≤ 0.1 and also when *b* is fixed but Δ*a, c* ≤ 0.1.

Turner99 is least stable for tRNA, while most stable is split between “best tRNA” and “best 5S.” In contrast, 5S rRNA is least stable for “best tRNA,” but most stable for “best 5S.” The “best both” are a good compromise, and certainly comparable to all Turner stabilities.

#### 3.1.3 Robustness

Stability analysis shows that small changes in multiloop initiation parameters, especially in *b*, may alter MFE predictions. We now investigate how different those predictions are.

Robustness *c* within error *r* means that even if the parameters are independently varied by ±*r*, the similarity of any new prediction to an original one is at least *c*. Robustness of all 6 parameters for both families is given in Tables 6 and 7 for *r* values that double starting with initial value .1.

Although MFE predictions are not necessarily stable within .1 error, the similarity remains high even as the parameters change. For both test families, the “best both” robustness is still greater than .5 at distance 0.4. For tRNA, this improves over Turner99 but is comparable to the other parameters. For 5S rRNA, this is no worse than “best 5S” or Turner89, and better than the other parameters.

### 3.2 Geometric details

RNA branching polytopes have certain distinctive characteristics. To determine if these are biologically meaningful, we compare the 50 real RNA sequences against two background distributions, the 50 shuffles and the 50 uniformly random ones, for each of the two test families.

Overall, there are some significant differences between the tRNA and 5S rRNA length scales. However, statistically significant differences within families did not occur with any consistent correlations that lead to biological insight.

#### 3.2.1 Polytope complexity

Computation time depends on polytope complexity, measured in terms of the number of vertices and of faces. Polytope complexity in turn depends on sequence length, as clearly seen in Table 8. For simplicity, computation time is reported in hours although it was measured in seconds.

An increase of less than 50 nt in average sequence length increases both the number of vertices and of faces by a factor of 3.6. This then increases the computation time from 2 hours to a full day. Beyond this, we can drawn no meaningful conclusion from the differences within the two families between the biological branching polytopes and those for the random sequences in terms of their complexity.

#### 3.2.2 Parameter space decomposition

Although an RNA branching polytope live in the 4d (*x, y, z, w*) coordinate space, we are only interested in the corresponding (*a, b, c, d*) parameters when *d* = 1. In this case, there is no scaling applied to the other NNTM values in the Δ*G* calculation. We also analyze the special case when *b* = 0, as the Turner99 and Turner04 parameters both use this value.

Each polytope still yields a subdivision of the 3d (*a, b, c*, 1) parameter space into connected, convex regions with flat sides. Now, though, the regions may be bounded as well as unbounded. The arrangement of unbounded regions in the (*a*, 0, *c*, 1) plane has a characteristic pattern, first illustrated in [32] and now fully described [34] for all fixed *b*.

Here we are concerned with the bounded regions since this includes the biologically realistic parameter ranges. The number of *d* = 1 regions under different constraints, bounded (bnd) and/or *b* = 0, is listed in Table 9. As with polytope complexity, ANOVA calculations did not identify any consistent, significant differences within the two families.

However, differences between families are again significant. For example, a greater percentage of 5S rRNA regions, 80 (2) versus 74 (3) for tRNA, intersect the *d* = 1 hyperplane. Of those, more 5S rRNA are bounded: 76% (1) versus 62% (3). The increase in sequence length increases the number of possible multiloops, which likely affects this distribution.

The fact that ∼50% of the polytope vertices may be of biological interest illustrates the challenge in improving prediction accuracy calculations. The numbers do drop substantially when *b* = 0, however all of the 3 “best” parameters identified here used *b* > 0.

The sensitivity of predictions to changes in *b* is explained by Table 10, which demonstrates that all regions are thin in *b*. The most significant differences are between families, although the lowest and highest values within families are different, but the overlap between families is not. A similar phenomenon happens in the *c* dimension, and also *a* when *x* > 1. It may be that these differences have biological implications, so we plan to investigate further in the future.

The high average *a* dimension is due to regions whose associated branching signatures have *x* = 1. When these regions are excluded, the average *a* length drops to roughly twice the *c* dimension. We do not understand the phenomenon yet, and plan to address it in a future study. Likewise, we will investigate the significant difference in the *a* value for the 5S rRNA sequences from all other test sets.

## 4 Conclusion

In this work we analyzed the effects of changing the three parameters (*a, b, c*) used in the initiation score, which approximates the entropic penalty, given to a multiloop in the NNTM. For this purpose we leveraged tools from geometry that allow us to build so-called branching polytopes for a diverse set of tRNA and 5S rRNA sequences and analyze all possible MFE structures for each of them. We then used this comprehensive information to give a complete analysis of the prediction accuracy, stability, and robustness for these sequences.

We find that on an individual basis, the secondary structure can be predicted with high accuracy (albeit never 100% accurately for 5S rRNA) for all sequences for some combination of multiloop parameters. This is a substantial improvement over the Turner99 parameters for a lot of sequences; however, the average maximum accuracy is not achievable for either tRNA or 5S rRNA for any choice of parameters. Using techniques from graph theory, we found combinations of parameters that improve the prediction for each family separately as well as across both families together. The “best both” parameters we found penalize the initiation of a multiloop more severely than the Turner99 parameters but then favor formations of branchings. We find that under these parameters the tRNA accuracy improves significantly whereas the difference in 5S rRNA accuracy versus the Turner parameters was not found to be significant.

Our analysis of the stability shows that the prediction is most sensitive in the change of the *b* parameter which is used to weight the unpaired nucleotides in the multiloops and least sensitive in the change of the parameter *a*. We explain this phenomenon by showing that the regions in the (*a, b, c*) parameter space that correspond to different predictions are significantly thinner in the *b* direction than in the other two. The robustness analysis shows that even though the prediction is not necessarily stable even under ±.1 error, the similarity of the predicted structures remains high even as the parameters change.

Finally, in order to determine whether the distinctive characteristics of the branching polytopes are biologically meaningful, we compared the complexity of the RNA branching polytopes to the one for two sets of random sequences: one set which was obtained by permuting the biological sequences in a way that preserves the dinucleotide frequency and one set in which the nucleotide frequencies are all equal to 25%. While some differences between the branching polytopes were observed, they were not significant enough to draw any meaningful conclusions. However, the complexity of the polytopes and the computational time needed for sequences at this length scale imply that the same kind of parametric analysis would be unfeasible for sequences of the order 1000 nt and to perform a similar analysis for such sequences a new algorithm would be needed.

## Acknowledgements

This work was supported by funds from the National Science Foundation (DMS 1815832 to SP and DMS 1815044 to CEH).

